# Clonal heterogeneity and antigenic stimulation shape persistence of the latent reservoir of HIV

**DOI:** 10.1101/2024.07.19.604385

**Authors:** Marco Garcia Noceda, John P. Barton

## Abstract

Drug treatment can control HIV-1 replication, but it cannot cure infection. This is because of a long-lived population of quiescent infected cells, known as the latent reservoir (LR), that can restart active replication even after decades of successful drug treatment. Many cells in the LR belong to highly expanded clones, but the processes underlying the clonal structure of the LR are unclear. Understanding the dynamics of the LR and the keys to its persistence is critical for developing an HIV-1 cure. Here we develop a quantitative model of LR dynamics that fits available patient data over time scales spanning from days to decades. We show that the interplay between antigenic stimulation and clonal heterogeneity shapes the dynamics of the LR. In particular, we find that large clones play a central role in long-term persistence, even though they rarely reactivate. Our results could inform the development of HIV-1 cure strategies.

## Introduction

Human immunodeficiency virus (HIV-1) actively replicates in CD4^+^ T cells ^1,2^. During the infection process, the genetic material of the virus is incorporated into the DNA of the host cell ^3^. Most infections result in the rapid production of new viruses, leading to the death of these infected cells within a few days ^4^. However, in a fraction of cells, HIV-1 is capable of lying in a dormant, “latent” state ^5^. This population of long-lived, latently infected cells is referred to as the latent reservoir (LR) of HIV-1. While antiretroviral therapy suppresses active HIV-1 replication, it is unable to eliminate latently infected cells or their integrated proviruses ^6^. Many latent viruses remain capable of reactivation, resulting in a quick return to active infection if antiretroviral treatment (ART) is interrupted, even in individuals who have undergone effective drug treatment for many years ^7,8^. The LR therefore presents the major barrier to an HIV-1 cure.

Understanding factors that contribute to LR persistence could greatly contribute to HIV-1 cure efforts. However, it is difficult to obtain a comprehensive picture of LR dynamics from direct measurements due to its small size. For typical HIV-1-infected individuals, roughly one in 10^*4*^ T cells are latently infected, and active HIV-1 replication occurs in only around 1% of these latently infected cells in viral outgrowth experiments ^9–11^. Thus, latently infected cells, especially ones that can readily reactivate, are rare. Subsequent studies have also found that multiple rounds of stimulation can prompt latent cells that initially remained dormant to reactivate, making it difficult to determine the total number of latently infected cells that are capable of reactivation ^11,12^.

Despite these challenges, recent work has provided in-sights into the dynamics and heterogeneity of the LR. Subsets of cells bearing integrated HIV-1 can undergo clonal expansion in patients receiving suppressive ART ^13,14^. The degree of expansion of clones as well as their persistence varies greatly and is associated with the specific integration sites ^13^ as well as stimulation by antigens ^15,16^. While it has been found that many highly-expanded clones contain defective proviruses ^17,18^, at least half of the cells carrying intact proviruses also belong to expanded clones ^18–21^. A strong negative correlation has also been observed between clone size and reactivation rate in viral outgrowth assays ^20^. As patients remain on ART for long times, the diversity of observed clones decreases and the proportion of HIV-1 proviruses in the largest clones progressively increases ^22^.

Mathematical modeling has also provided insights into the LR, with some model predictions validated in experiments ^23^. Studies investigating the relationship between latently infected cells and plasma viremia during ART ^24–29^ suggest that as long as ART is marginally effective, the persistence of latent virus is most strongly influenced by the longevity of infected cells and the rate at which they reactivate. Recent modeling work has also suggested that uneven homeostatic proliferation of latently infected cells early in infection may lead to the observed spread in clone sizes in the LR ^30^.

To gain a deeper understanding of the persistence of the latent reservoir (LR) in HIV-infected individuals, it is important to develop mathematical models that reflect the biological mechanisms that govern its dynamics. The LR is composed of diverse clones with different T cell receptors (TCRs), which affect their activation potential and antigen specificity. Moreover, these clones have distinct viral integration sites, which influence their transcriptional activity and reactivation probability. These factors contribute to the clonal heterogeneity of the LR and its persistence. Thus, there is a need to incorporate more comprehensive and biologically motivated features of clonal heterogeneity, which are not typically incorporated in existing mathematical models of the LR.

We addressed this challenge by developing a novel stochastic model of LR dynamics that explicitly accounts for clonal heterogeneity. We consider genetic changes in HIV-1 sequences and variable probabilities of reactivation, while also incorporating the effects of antigenic stimulation on latently infected clones with different T cell receptors. The dynamics of these clonal populations are integrated with interactions between free viruses, susceptible cells, and cells that are actively infected.

Our model recapitulates experimentally observed features of HIV-1 infection while also providing insights into LR structure, dynamics, and persistence. We recover the decay kinetics of HIV-1 RNA in blood, HIV-1 DNA in peripheral blood mononuclear cells (PBMCs), and latent cells that reactivate upon stimulation, which occur over widely-varying time scales (days, months, and years) following the start of ART ^31–33^, without the use of time-varying parameters. Among other findings, our model reproduces the observation of defective proviruses in highly expanded clones ^17^ and the negative correlation found between clone size and reactivation probability for patients who have undergone ART treatment for many years ^20^. We find that stimulation by antigens combined with heterogeneous reactivation rates for different clones leads to a broad distribution of clone sizes, which are stratified by their reactivation rates. Over long times, we find that the LR becomes progressively more concentrated on a small number of clones with low reactivation rates, which play a key role in LR persistence. These insights could inform the development of new therapeutic approaches to reduce the size of the LR and achieve a functional HIV-1 cure.

## Results

### Stochastic model incorporating LR heterogeneity

Inspired by past modeling work ^31,34–36^, our model (**Figure 1A**) consists of four main populations: uninfected CD4^+^ T cells (T), productively infected activated CD4^+^ T cells (A), latently infected resting CD4^+^ T cells (L), and HIV-1 virions (V). All cells have finite lifespans determined by their respective death rates (see Methods for a complete list of parameters and supporting references). We used a constant replacement rate λ_*T*_ to approximate the replenishment of uninfected target cells from the thymus ^37–39^. The rate of production of virions is given by the product of the death rate of actively infected cells *µ*_*A*_ ^40^ and the viral burst size *n* ^41^. Virions are then cleared at a constant rate *c* ^42^.

**Fig. 1.**
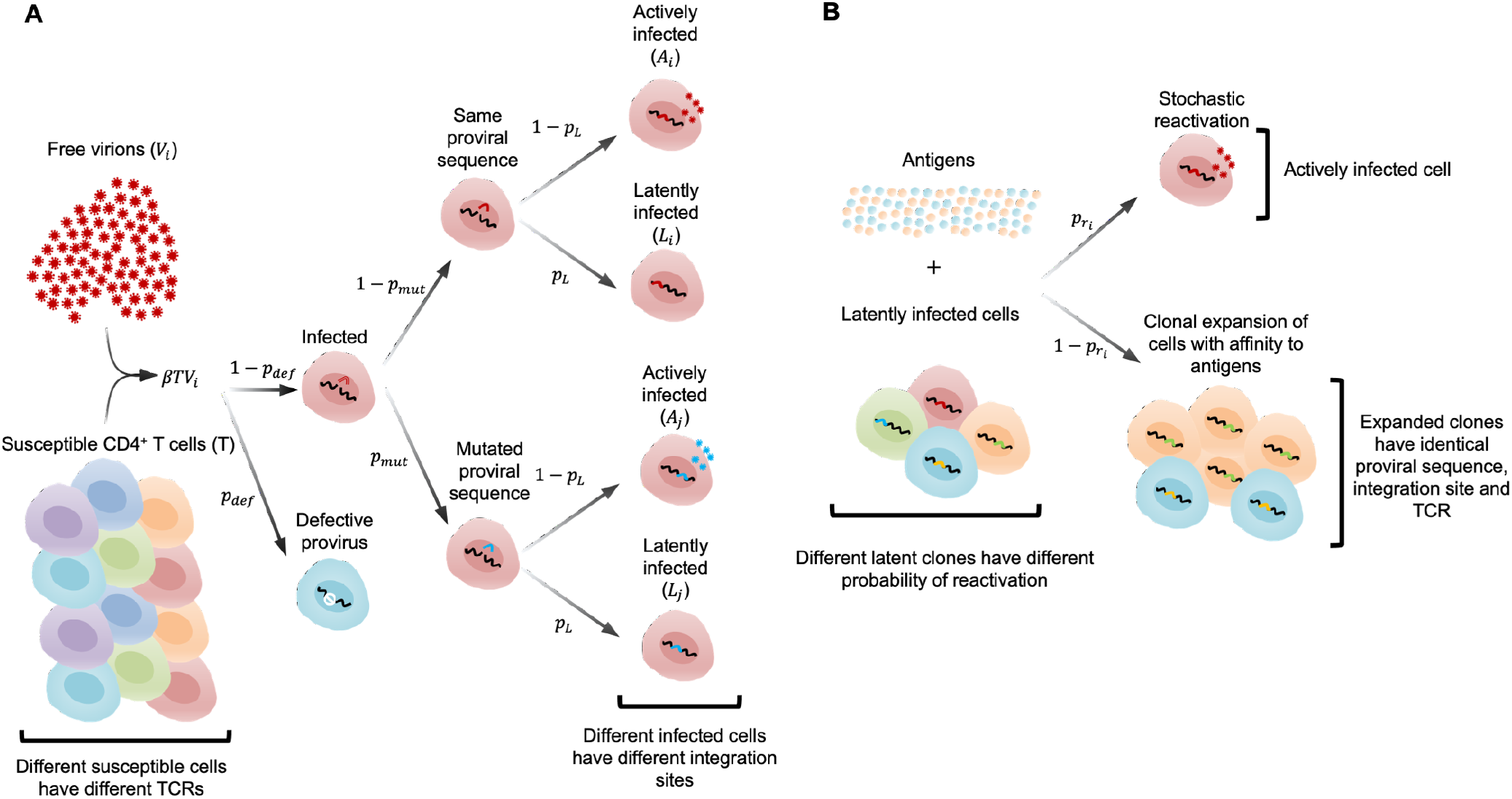
Model schematic. (**A**) When a new cell is infected, it forms a new “clone” with a specific TCR sequence. The generation of defective proviruses, point mutations, and active or latent infection are all determined by chance. (**B**) Clonal expansion and reactivation are possible outcomes of the dynamics of latently infected clones upon stimulation by antigens.

HIV-1 virions can infect susceptible CD4^+^ T cells. A small fraction of infections, *p*_def_, will result in defective integrated proviruses due to effects like large deletions or hypermutation. During ART, this leads to proviruses with fatal defects outnumbering intact proviruses by a factor of 10−50 to 1^*22*^. HIV-1 also mutates during infection due to error-prone reverse transcription, with an estimated error rate of 3 × 10^*−*5^ per base per replication cycle ^43–45^. Given the length of the HIV-1 genome, this implies that approximately a third (*p*_mut_) of successful infection events will lead to a mutation. Finally, a fraction *p*_L_ of infection events will result in latent rather than active infection. We fit *p*_L_ such that the HIV-1 DNA per 10^*6*^ PBMCs at the beginning of ART was in the range *−*10^*3*^ 10^*4*^, consistent with patient data ^31^. Consistent with reported HIV-1 RNA levels during ART ^32,46^ and following previous modeling studies ^28^, we assume that viral replication is attenuated but not perfectly suppressed during ART.

To account for the heterogeneity of latently infected cells, we modeled each latently infected clone individually, including the expansion and antigen-driven proliferation of individual clones (**Figure 1B**). Clones are defined as latently infected cells with identical T cell receptors, integrated proviruses, and integration sites. Each time a new clone is created through a latent infection event, it is assigned a random probability of reactivation (Methods), consistent with the observation that integration is stochastic and different integration sites can affect the capacity for reactivation ^47^. In addition, each clone is stimulated by a background concentration of antigen that fluctuates over time (Methods), inspired by past models of T cell repertoire dynamics ^48^. We assume that the homeostatic death and proliferation rates are the same for all clones; however, stochastic differences in antigenic stimulation drive differences in clonal proliferation. Here our approach differs from previous models that considered a constant rate of reactivation for latently infected cells ^28^ or different cell populations with different half-lives ^27^. We simulated our model using a system of stochastic differential equations that describe the dynamics of cells during HIV-1 infection both before and during ART (Methods).

### Seeding of the reservoir and clonal proliferation pre-ART

We first simulated the seeding and development of the latent reservoir during the first months of infection. Simulations begin with a number of virions in the system and zero active and latently infected cells (Methods). Even during the first weeks of infection, we observed a large number of distinct latently infected clones in the reservoir. Most clones are very small: for up to a year after infection, the average clone size is less than 10 cells, with a median clone size of 2 cells (**Figure 2**). During this time, the largest clone is typically smaller than 1000 cells. Most of these clones are also shortlived. The average age of a clone after one year of infection is 15 days, with a median age of 6 days. Such short-lived clones are highly likely to reactivate, with an average probability of reactivation *p*_*R*_ around 9%.

**Fig. 2.**
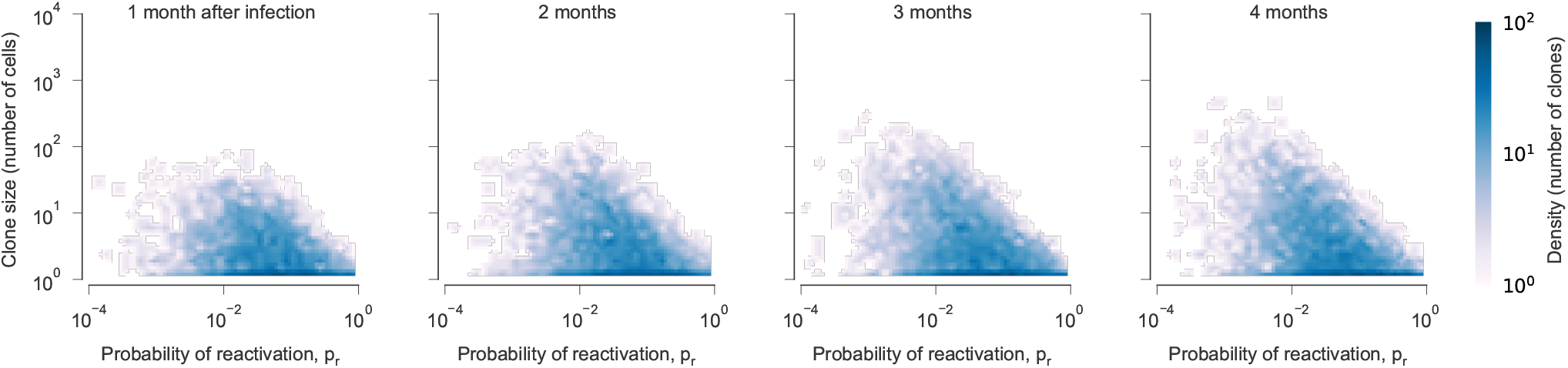
Distribution of clones in the HIV-1 latent reservoir early in infection. Most clones are small and have high reactivation probabilities. However, as time passes, clones with lower reactivation probabilities begin to grow in size. Stimulation from antigens drives some rare clones with very low probabilities of reactivation to large sizes.

During this early phase, some clones will be stimulated to proliferate by exposure to antigens. However, the effect on different clones in the reservoir differs substantially depending on how likely the latent virus is to reactivate when stimulated. In clones with low reactivation probability, proliferation due to antigenic stimulation typically results in net growth. In clones that readily reactivate, however, the reactivation of latent virus dominates, suppressing proliferation or even leading to the elimination of these clones. These dynamics lead to a progressive increase in the number of clones with low reactivation probabilities and large clone sizes over time.

### Kinetics of plasma viral load, HIV-1 DNA, and the inducible viral reservoir after ART initiation

After the LR has been seeded, we simulated the response of viral populations, including both latent and actively infected cells, to long-term ART. To simulate viral kinetics during ART, we decreased viral infectivity *β* due to treatment (Methods), keeping all other parameters constant.

Clinical data shows that after initiating ART, HIV-1 RNA in blood decreases rapidly over the course of around 2 weeks, with observed half-lives 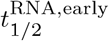 of 0.9-1.9 days ^31,50^. This is followed by a more gradual decline over the next 4 weeks. 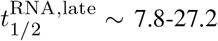days ^31,50^ The total number of latently infected cells, measured by HIV-1 DNA in PBMCs, decays steadily over the first few months on ART 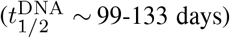. Infectious units per million PBMCs (IUPM), measured in viral outgrowth assays (Methods) declines very slowly, with a measured half-life 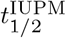 of approximately 44 months ^49^.

Our model quantitatively recovers the decay rates of HIV1 RNA and latently infected cells spanning days, months, and years on ART (**Figure 3**). In our simulations, viral load first drops sharply, which is primarily driven by the death of actively infected cells (**Figure 3A**). At the same time, small clones are gradually eliminated through reactivation or random cell death. Due to reduced viral replication, these clones are no longer replenished at the same rate, leading to a net decline in the total number of latently infected cells. Clones with high reactivation probabilities are depleted more rapidly than those that do not readily reactivate. Collectively, these factors lead to a shift in the LR toward larger clones with lower rates of reactivation, slowing the decline in viral load and HIV-1 DNA in PBMCs (**Figure 3A-B**). As larger clones are slowly eliminated, we find a decline in the inducible reservoir consistent with measurements from clinical data (**Figure 3C**).

**Fig. 3.**
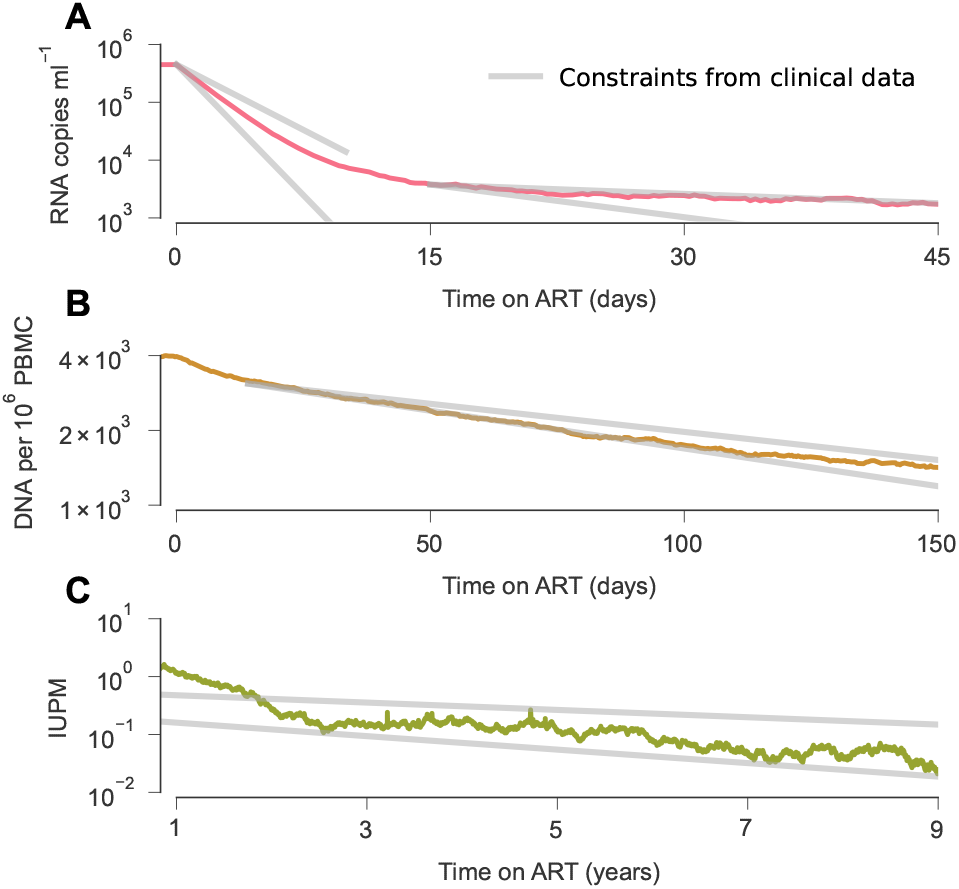
Latent reservoir dynamics after ART initiation in an example simulation. (**A**) ART first results in the rapid decline of plasma viral load ^31^, first due to the death of actively infected cells and later driven by the elimination of small clones in the LR. (**B**) Over slightly longer times, the number of latently infected cells steadily declines ^31^ as small clones die out and are no longer quickly replenished by new infections. (**C**) Infectious units per million (IUPM), a measure of cells in the LR capable of reactivation, declines over the course of years, approximately following the 44-month half-life measured in clinical data ^49^.

### Long-term clonal dynamics in the latent reservoir

During ART, clones with higher probabilities of reactivation have a shorter effective survival time than clones with lower probabilities of reactivation. We therefore find that the average probability of reactivation decays over time. However, clones that readily reactivate are not entirely eliminated. Occasional reactivation of latent cells from large clones leads to bursts of viral replication that partially reseed the reservoir. These dynamics result in a long-term quasi-steady state, where small clones with high probabilities of reactivation turn over frequently while large, quiescent clones slowly fluctuate in frequency.

Over long times, we find that, for the largest clones, clone size *n* scales inversely with the probability of reactivation 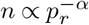 with an exponent *α* ∼ 1 (**Figure 4A**). This finding is consistent with previous work that observed a power law relationship between clone size and probability of reactivation in viral outgrowth assays in data from multiple subjects years after ART initiation ^20^. Clones that are small and/or have low probabilities of reactivation (i.e., ones occupying the lower left corners in **Figure 4A**) are particularly challenging to quantify in patient data because their probabilities of being sampled in sequencing HIV-1 DNA from PBMCs or viral outgrowth assays are exceedingly small. Such clones are likely to be observed only once in data if they are sampled, which is consistent with observations in clinical data ^20^.

**Fig. 4.**
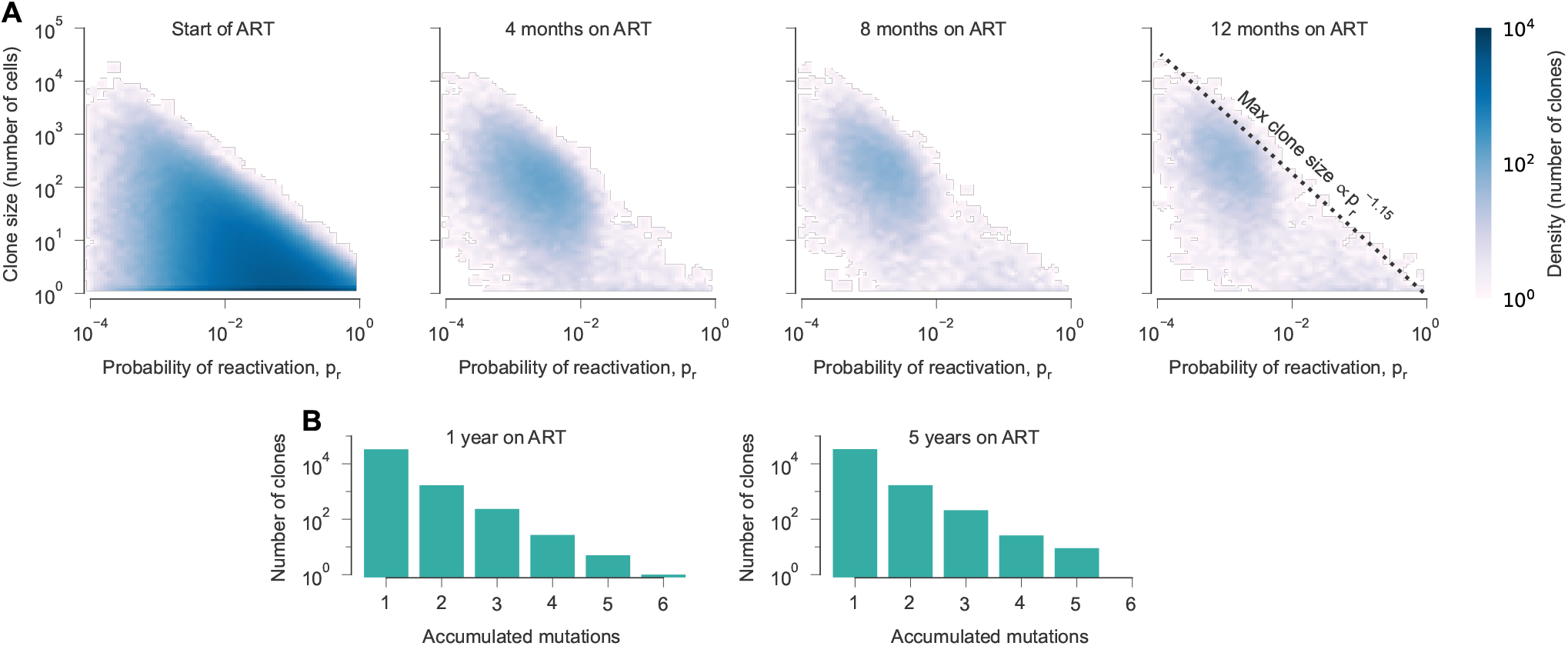
Dynamics of the latent reservoir during ART. (**A**) As ART begins, many clones are observed with different sizes and probabilities of reactivation. Reactivation and random fluctuations lead to the preferential loss of small clones and ones with high probabilities of activation over time. The largest clone size scales roughly with the inverse of the reactivation probability, a relationship that is stable over time. (**B**) Despite viral replication, few mutations accumulate in latent clones during ART.

Our results also show that after a year on ART, the relative size of large clones changes little over time. Collectively, our simulations are consistent with longitudinal studies that found few significant changes in the proportion of different clones in the LR when sequencing proviral DNA ^22,51,52^, while significant changes in clonal distributions can be observed when sequencing reactivated viruses from viral outgrowth assays ^51,52^.

### Presence or absence of HIV-1 evolution during ART

While ART strongly suppresses viral replication, it may not be completely effective. Past modeling work has suggested that low amounts of replication can continue after treatment intensification and may influence the level of detectable virus, but are unlikely to allow for long-term sequence evolution ^25,27–29^. Others have argued that ongoing HIV-1 replication can lead to measurable viral evolution during ART ^53–55^, though this point is hotly debated ^56,57^, and multiple studies have failed to observe evolution in the reservoir during ART^58–61^.

To test whether or not viral sequence evolution would occur in our model, we tracked the number of accumulated mutations in individual clones after ART initiation. In our simulations, the mean number of new infection events resulting from active infection in a single cell is smaller than 1 due to the suppressive effects of ART. This implies that persistent, self-sustaining active replication is impossible. However, because our model is stochastic, we observe occasional “bursts” of viral replication. Despite occasional bursts, we found no evidence for progressive accumulation of mutations or genetic divergence over time (**Figure 4B**), consistent with past observations ^58–61^ and modeling work ^25,27–29^.

One prominent prior study that argued for continued HIV-1 evolution during ART was based on observations made during the first year of ART ^55^. As stated above, we do not find evidence for significant sequence evolution in our model. However, we do observe immense changes in individual clone sizes during early ART, especially for many small clones that are eliminated. These dynamics support previous arguments that sampling of different clones could explain the *appearance* of evolution shortly following ART ^62^.

### Effects of early intervention or elite control on LR structure

Recent studies found that the composition of the LR is altered in individuals who have undergone early ART treatment ^63^ and in elite controllers, who naturally maintain very low levels of viral replication even in the absence of ART ^64^. To mimic elite control or early ART, we adjusted our simulations to include a sharp drop in viral infectivity 12 days after initial HIV-1 infection (Methods). Unlike previous simulations, early control results in a much smaller number of clones in the LR (**Figure 5A**), which quickly becomes dominated by just a few large clones (**Figure 5B**). Our simulations thus recapitulate studies showing that the LR in elite controllers or those with early ART are mono-or oligo-clonal, with little reactivation and background replication ^63,64^.

**Fig. 5.**
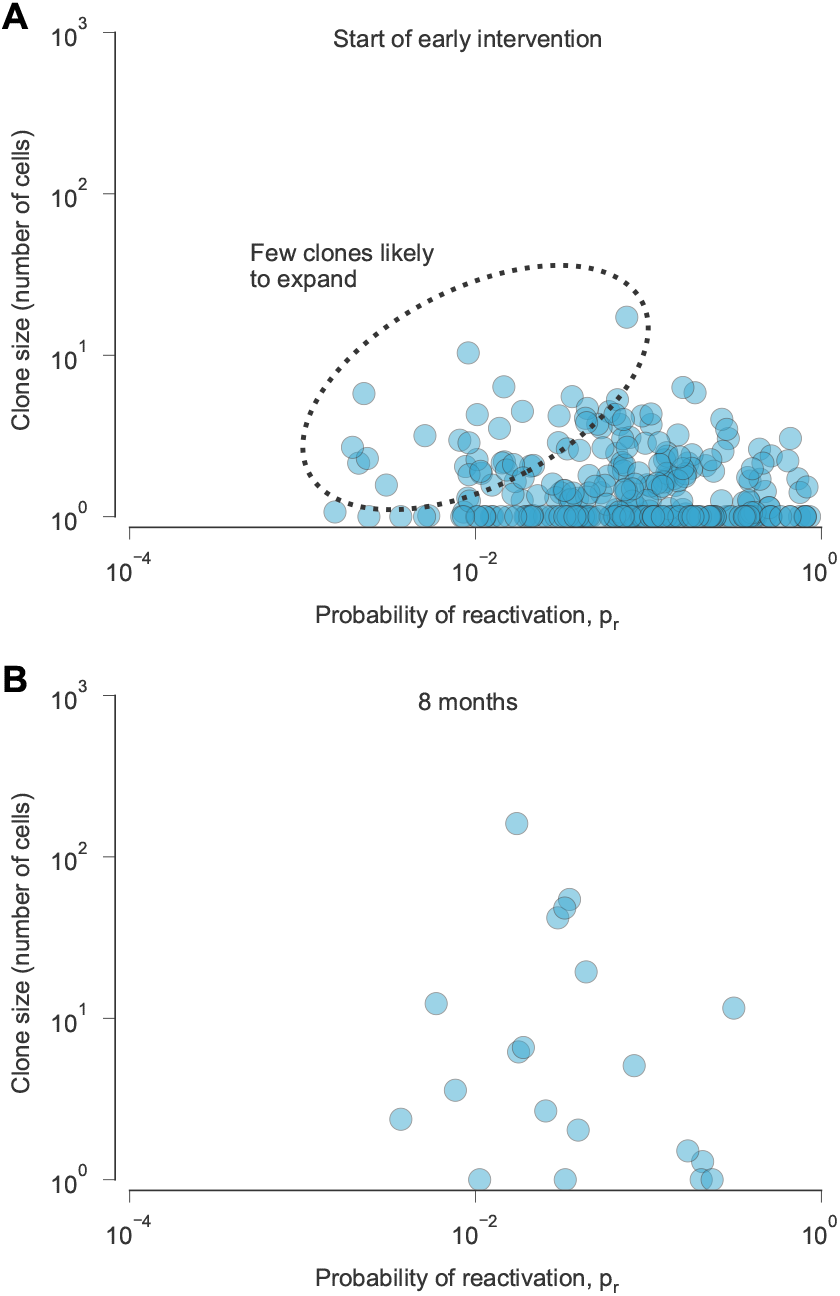
Distribution of clones in the latent reservoir after early intervention. **(A)**We simulated the effects of early intervention by initiating ART conditions 12 days after infection. At this time, there are few clones that are large or have small probabilities of reactivation, limiting the potential for clonal expansion. (**B**) After 8 months, few clones remain.

### Factors underlying long-term LR persistence

Despite suppressive ART, the latent reservoir persists for decades in HIV-1-infected individuals. How do different components of the LR contribute to its persistence? We find that the presence of large latent clones is the most important factor in the long-term persistence of the latent reservoir, despite their low probabilities of reactivation. In fact, small *p*_*r*_ allows these clones to grow to large sizes with minimal decay due to reactivation (**Figure 4A**). Due to their size, these clones are also very unlikely to die stochastically due to fluctuations in clone size, unlike smaller clones that turn over rapidly.

Interestingly, the largest clones are not the ones that are most likely to generate rebound viruses. On average, the rate of viral outgrowth is proportional to the product of the clone size and reactivation probability. In typical simulations, this is maximized by clones with intermediate sizes and reactivation probabilities. This is because clones with very small probabilities of reactivation are fairly rare, and clones with very high probabilities of reactivation tend to be small and short-lived. Thus, even though we find that the largest clones (with small probabilities of reactivation) are chiefly responsible for LR persistence, they may not be the typical first source of outgrowing viruses during rebound.

## Discussion

Here we developed a stochastic model of the latent reservoir of HIV-1 that accounts for the inherent heterogeneity of different clones in the reservoir. Our mechanistic approach and direct simulation of each clone allows us to delve deeper into the underlying processes shaping reservoir dynamics. Our model recapitulates changes in HIV-1 measurements in clinical data over the scale of days (decline in HIV-1 RNA in the blood after ART) to years (decline in IUPM over years on ART). We also quantitatively recover a “power law” relationship between clone size and reactivation for large clones, and we find realistic distributions of clones in the LR in simulations that mimic early intervention with antiretroviral drug treatment.

Beyond comparisons with experimental and clinical data, our model makes several predictions about the structure and long-term dynamics of the latent reservoir. Our study suggests that the LR consists of a very large number of clones, especially small clones. Due to their small size and (for some clones) low probability of reactivation when stimulated, they would be difficult to detect through conventional means in studies that seek to characterize the viral reservoir. Nonetheless, collectively, they contribute to the diversity of the latent reservoir and serve as a potential source of viral rebound.

Clonal heterogeneity, including the propensity of different clones for reactivation and stimulation by antigens, emerged as a critical factor to reconcile both short- and long-term dynamics of the latent reservoir after ART. Differences in probabilities of reactivation lead to a slow but progressive “coarsening” of the reservoir, as clones that readily reactivate are eliminated and larger, quiescent ones persist. The persistence of clones with low probabilities of reactivation in our model aligns with recent longitudinal studies that have reported the positive selection of proviruses with lower transcriptional activity during prolonged ART ^65^.

Our simulations also show nuanced effects of sporadic viral replication during ART. With zero viral replication, all small clones in the LR would ultimately be eliminated due to random clone size fluctuations. However, the level of active replication needed to sustain a population of small clones is insufficient to produce long-term sequence evolution of the virus ^25,27–29^. This emphasizes an important distinction between viral replication and evolution, especially evolution within the LR. Persistent, self-sustaining viral replication will lead to the accumulation of mutations (i.e., evolution) over time. However, the same is not true for sporadic bursts of replication that cannot be sustained, and which must be restarted from the same pool of unmutated latent viruses after previous active infections die out. Our model suggests that sporadic replication during ART is consistent with experimental data, but persistent replication is not.

Past work has identified various factors that could affect clonal proliferation, including different HIV-1 integration sites ^47^. Here, we found that random differences in antigenic stimulation alone are sufficient to reproduce the observed structure of the latent reservoir. This result should not be interpreted as evidence that different integration sites do not play a role in heterogeneous clonal expansion. Rather, our work shows that differences in antigenic stimulation can already lead to stratification in clone sizes and dynamics. Additional factors could also further contribute to the heterogeneity of the LR. For example, a proliferative advantage associated with specific integration sites could promote clonal expansion, potentially extending the lifetime of the reservoir.

Many assays have been devised to quantify the magnitude and diversity of the HIV-1 latent reservoir, but quantifying the true size of the LR remains challenging ^11^. To address this, a hybrid approach combining stochastic modeling and statistical analysis that accounts for the limitations of experimental noise, similar to proposals for T cell repertoire diversity estimation ^66^, may offer an effective quantitative measurement of the LR. Our work could contribute to this effort by providing a way to study small clones, which are difficult to access experimentally, using a rigorous model constrained by experimental and clinical data.

Understanding the structure and dynamics of the HIV-1 latent reservoir could aid in the development of HIV-1 cure strategies that aim to eliminate or permanently suppress the LR. Our model contributes to these efforts by providing a quantitative description of the LR that is consistent with existing data, but which also extends to “unseen” areas that are difficult to characterize experimentally. One important finding relevant for HIV-1 cure strategies is that large clones that are replication-competent but relatively unlikely to reactivate play a key role in long-term persistence of the LR. In future work, our model could be used to simulate responses to different types of therapeutic interventions, evaluating plausible paths to an HIV-1 cure.

## ACKNOWLEDGEMENTS

Computations were performed using the computer clusters and data storage resources of the University of California, Riverside High Performance Computing Cluster (UCR HPCC). The UCR HPCC is funded in part by grants from NSF (MRI-1429826) and NIH (1S10OD016290-01A1). The authors declare no competing interests.

## AUTHOR CONTRIBUTIONS

Author contributions: M.G.N. and J.P.B. designed research; M.G.N. performed research; M.G.N. and J.P.B. analyzed data and wrote the paper.

## Methods

### Mathematical model

Our model consists of four main populations: uninfected CD4^+^ T cells (T), productively infected activated CD4^+^ T cells (A), latently infected resting CD4^+^ T cells (L), and HIV-1 virions (V). All cells have finite lifespans determined by their respective death rates, and virions are cleared at a clearance rate *c*. Uninfected cells are produced at a constant, fixed rate, and virions at a rate proportional to the number of active cells. HIV-1 virions can infect target CD4^+^ T cells, and upon infection, a small fraction will result in defective cells due to effects like large deletions or hyper-mutations. Functional proviruses will accumulate a mutation with probability *p*_mut_. Finally, a small fraction of infection events will result in latently infected cells. To account for stochasticity, these processes are modeled with probabilities instead of rates.

To account for the heterogeneity of the LR in our model, we define a latently infected clone as a set of cells that have the same TCR, proviral DNA sequence, and integration site. Individual clones will differ in how they are stimulated by antigens and their propensity for reactivation. Due to differences in integration sites, each new clone *L*_*i*_ is assigned a random probability of reactivation 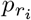. Following work describing dynamics of T cell repertoires ^67^, we describe the stimulation of each clone by antigens with 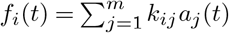 where *k*_*ij*_ is the interaction coefficient between clone *i* and antigen *j* (when clones are cross-reactive), and 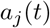 is the overall concentration of an antigen *j* as a function of time. We assume that antigen concentration decays exponentially after its introduction at random times as pathogens are encountered and cleared, either passively or through the action of the immune response.

When a latently infected cell is stimulated to divide, there is a probability of the latent provirus reactivating, converting the cell into an actively infected cell. In this case, the number of latent cells decreases by 1 and the number of actively infected cells increases by 1. If no reactivation occurs, then the latently infected cell proceeds to divide in response to the antigen interaction.

The dynamics followed by a latently infected clone *L*_*i*_ are then driven by its basal division rate *v*_*L*_, death rate *µ*_*L*_, probability of reactivation 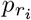, and its interaction with antigens *f*_*i*_(*t*). This gives the following stochastic differential equation (SDE):

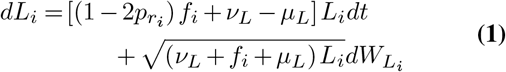

As mentioned above, the function *f*_*i*_(*t*) encodes the fluctuations of the environments as experienced by clone *i*. The stochastic process giving rise to *f*_*i*_(*t*) is a sum of Poisson-distributed, exponentially decaying spikes. This process is not easily amenable to analytical treatment or simulations, so following the approach of Desponds et al. ^67^, we assume that correlations among clones are weak and replace the function with a simpler one with the same temporal autocorrelation, that is an Ornstein-Uhlenbeck process:

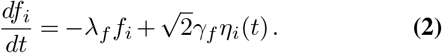

Here *η*_*i*_(*t*) is a Gaussian white noise, λ_*f*_ is the inverse of the characteristic lifetime of antigens, and *γ*_*f*_ quantifies the strength of variability of the antigenic environment.

To model the active cells and virions we need to consider what happens during an infection event, which is illustrated in **Figure 1**. A virion with sequence *k* finds and successfully infects a susceptible T cell at rate *β*. During infection, there is a probability *p*_def_ that the integrated provirus will be defective due to large deletions, hypermutation, or other similar alterations. For proviruses that are not defective, we model the accumulation of point mutations with probability *p*_mut_. Finally, we consider a probability *p*_*L*_ for the infection to be latent. Each latent infection defines a new latent clone, since we assume that the probability that two identical viruses integrate at the same location in two T cells with identical T cell receptors in separate infection events is essentially zero. This clone could share the same sequence as another latently infected clone but have a very different integration site and therefore a different probability of reactivation.

Collectively, the dynamics governing actively infected cells and virions are defined by the following system of SDEs:

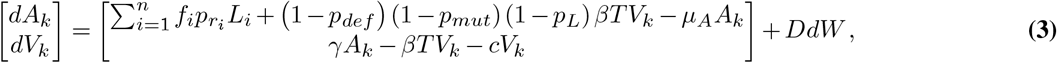

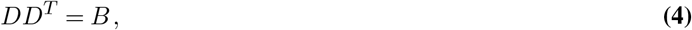

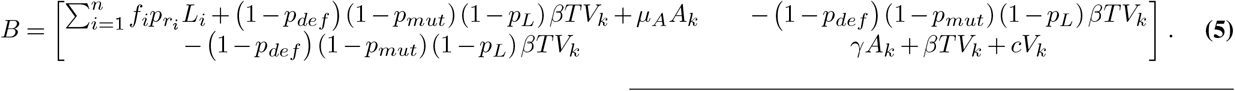

Finally, the susceptible T cells in our model follow the simple stochastic differential equation

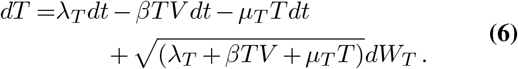

### Modeling the reactivation probability distribution

Evidence suggests that different latent clones have different propensities for reactivation ^68–72^. Ultimately, this heterogeneity in reactivation plays a central role in concisely explaining widely differing and multiphasic decay rates of HIV-1 RNA in blood, HIV-1 DNA in PBMCs, and IUPM observed in clinical data. Of course, heterogeneous reactivation rates are also essential to reproduce the observed association between the probability of reactivation in viral outgrowth assays and latent clone size.

However, there is no singular distribution of reactivation probabilities that can uniquely reproduce the observed clinical values. For simplicity, we opted to use a lognormal distribution, which is a natural choice when a variable (i.e., probability of reactivation) is obtained as the product of many independent explanatory variables (e.g., virus genetic background, cell type, integration site, local chromatin context, etc. ^72^). Previous modeling work has used a lognormal distribution for turnover rates to define uneven proliferation of clones in the first year of infection ^73^. Modeling the probabilities of reactivation with this distribution, which is grounded in the underlying biology of latent infection and consistent with clinical data, allows us to extend our model not only to active infection but also to describe long-term dynamics of the latent reservoir during ART and viral rebound.

We explored various values of *µ*, the average of the logarithm of probabilities of reactivation. We then adjusted *σ*, the spread of the logarithm of probabilities of reactivation to align with the multiphasic decay patterns observed in viral load, HIV-1 DNA, and IUPM measurements. Generally, as *µ* increases, the number of clones with high probabilities of reactivation and smaller sizes increases. This leads to a sharper drop in HIV-1 DNA post ART and a shorter time for the reservoir to become oligo-/monoclonal. Thus, the values of *µ* and *σ* are constrained, if not completely determined, by existing clinical data. To illustrate our findings, we used a log-normal distribution with *µ* = −1 and *σ* = 0.8. Experimental measurements of the distribution of probabilities of reactivation would be of great interest, allowing us to more precisely model long-term behavior of the reservoir.

### Modeling seeding of the reservoir and ART initiation

To simulate the initial establishment of the reservoir, we first calibrated the parameter *β* to capture the observed rise in viral load during the first two Fiebig stages of infection ^74,75^. Subsequently, after a month, we decreased the value of *β* to emulate the immune system’s suppression of viral replication, while maintaining fixed conditions for clonal proliferation prior to ART initiation. This adjusted*β* value corresponds to the viral load levels observed during stages 4 and 5, and these conditions remain constant until the initiation of ART, which in our primary example simulation occurs 60 months post-infection.

Upon ART initiation, the impact of treatment is simulated by modifying the value of *β*. Specifically, this adjustment aims to align the initial decline in viral load in simulations with the decay of HIV-1 RNA in blood observed in clinical data during the first two weeks following ART initiation ^76^. Given that the number of virions we observe is proportional to the number of active cells and to the viral load, we follow the methodology outlined in Hill et al. ^77^ to estimate viral load, where the number of actively infected cells is divided by 1680. This value is the geometric mean of different estimates from clinical data for the cell to virus ratio, obtained by balancing viral production and decay at equilibrium with an estimate that virus particles in the lymphoid tissue outnumber the ones in circulation 100-fold. Throughout the pre-ART and ART periods, all other parameters are held constant and remain unchanged.

To replicate scenarios involving elite control or early ART initiation, we introduced a rapid decline in infectivity (*β*) shortly after the initial HIV-1 infection. Rather than waiting for 60 months to commence ART, we transitioned to the *β* _*ART*_ value after only half a month of infection.

### Metrics for quantifying the HIV-1 latent reservoir and infection

We used infectious units per million (IUPM) to quantify the abundance of replication-competent HIV-1, which is measured in viral outgrowth assays. In our simulations, we used the product of each non-defective clone’s size and its probability of reactivation, summed over all clones, as a proxy for IUPM. This quantity should indeed be proportional to the probability that a latent clone is sampled and successfully reactivates when stimulated, which is analogous to IUPM.

We quantified HIV-1 DNA per 10^*6*^ peripheral blood mononuclear cells (PBMCs) by dividing the total number of latent cells by the total number of T cells and multiplying the result by 10^*6*^.

As described above, we quantified HIV-1 RNA in blood in our simulations by dividing the current number of actively infected cells by 1680, the geometric mean of different estimates for the cell to virus ratio, obtained by balancing viral production and decay at equilibrium with an estimate that virus particles in the lymphoid tissue outnumber the ones in circulation 100-fold ^77^.

### Computational implementation

We used the Euler-Maruyama method to simulate the dynamics of our system of stochastic differential equations (SDEs) ^78^. This numerical technique allows us to approximate the deterministic component of the SDEs using the Euler method at each time step. To incorporate the stochastic component, a random term is introduced, generated by a normally distributed random number with a mean of zero. The standard deviation of this random term was determined by the coefficients present in the SDEs.

Due to the complexity of our model, it was not computationally feasible to simulate the full model using a realistic number of CD4^+^ T cells, roughly 1.75 × 10^*11*^ for a typical adult ^79^. We therefore used two complementary approaches to perform our simulations. First, we simulated the full model at smaller system sizes (i.e., numbers of CD4^+^ T cells). Second, we developed and simulated simplified models that could readily scale to larger system sizes. As described in sections below, we carefully compared the output of both the full and simplified models for smaller system sizes to ensure that the simplified models accurately captured LR dynamics from the full model.

Here we refer to the order of magnitude of a simulation as the total number of CD4^+^ T cells included in the simulation. For example, we refer to a simulation including a realistic number of CD4^+^ T cells for an adult, around 1.75 × 10^*11*^, as a simulation at order 11. The figures presented in the full simulation were based on an order of magnitude of 7, that is, a total number of CD4^+^ T cells of 1.75 × 10^*7*^. In these simulations, the thymic production of T cells, viral infectivity, and metrics for quantifying HIV-1 presence and the LR are scaled in proportion to the total number of T cells in the simulation. For example, if the total number of CD4^+^ T cells decreases by an order of magnitude, the infectivity *β* increases an order of magnitude such that the product *β T* remains the same.

### Simplified model for latent clones

It is not possible to make full simulations at higher orders of magnitude than 7 due to computational limitations. This order of magnitude however does not allow us to fully appreciate changes in the clone size distribution during ART as the majority of clones die. To overcome this, we use a simplified model for latent clones alone, assuming a constant influx of new clones during active infection and a different constant rate of new clones during ART. These values were calibrated based on fully detailed simulations at order 7 such that the decays of latent cells, number of clones, and changes in the distribution matched between the full simulation and the simplified one (**Supplementary Figs. 1** and **2**). These parameter values were then scaled up to run the simplified simulation and make the figure shown in the main text for dynamics of the latent reservoir during the first year on ART.

### Simplified model for tracking mutations

We further extended our simplified model to study the number of accumulated mutations during ART. To develop this model, we assumed a constant rate of mutation accumulation, derived from the full simulation at order 7 and rescaled as described above. Conservatively, we used the maximum number of new mutant latent clones and active infections produced per time step over a window of one month after one year on ART (**Supplementary Fig. 3A**). Because these mutation accumulation rates were based on the maximum rate of mutation accumulation in full simulations, we expect that the number of mutations accumulated in our simplified simulation should serve as a conservative upper bound on the true number of accumulated mutations in a complete simulation.

## Data and code

Raw data and code used in our analysis is available in the GitHub repository located at https://github.com/bartonlab/paper-HIV-latent-reservoir. This repository also contains Jupyter notebooks that can be run to reproduce the results presented here.

---

**Table 1.** Table of parameters used in the model.

| Parameter | Value | Description | Type | Source |
| --- | --- | --- | --- | --- |
| $p_{\text{def}}$ | 0.002 | Probability that an infection event results in a defective HIV-1 provirus. | Fitted | Fitted to recover ratios between defective and intact proviruses during ART of 10 – 50 to 1 (ref. <sup>80</sup>). |
| $p_{\text{mut}}$ | 0.33 | Probability of a mutation during each infection event. | Literature | Obtained from estimates of the average HIV-1 mutation rate and genome length <sup>81–83</sup>. |
| $p_L$ | 0.05 | Probability that an infection event will result in a latent rather than active infection. | Fitted | Fitted to recover total levels of HIV-1 DNA per $10^6$ PBMC at the beginning of ART measured in patient data <sup>76</sup>. |
| $\lambda_T$ | $1.05 \times 10^{10} \text{ cells month}^{-1}$ | Replacement rate of uninfected target cells from the thymus. | Literature | $3.5 \times 10^8$ CD4<sup>+</sup> T cells per day at age 20, converted to months <sup>84</sup>. |
| $\mu_T$ | $0.06 \text{ month}^{-1}$ | Net decay of susceptible cells. Equivalent to the difference between proliferation and homeostatic death of susceptible cells. | Literature | Value needed to maintain typical size of the CD4<sup>+</sup> T cell compartment <sup>79</sup>. |
| $\beta_{EI}$ | $4.0 \times 10^{-13}$ | Infectivity rate during the first month of infection. | Fitted | Fitted to the rate of viral load increase observed during the first 2 Fiebig stages <sup>75</sup>. |
| $\beta_{AI}$ | $1.8 \times 10^{-13}$ | Infectivity rate during active infection post immune response. | Fitted | Fitted to recover the viral load observed during Fiebig stages 4 and 5 (ref. <sup>75</sup>). |
| $\beta_{ART}$ | $6 \times 10^{-14}$ | Infectivity rate during ART. | Fitted | Fitted to recover the first stage of viral load decay post ART <sup>76</sup>. |
| $\mu_A$ | $21 \text{ month}^{-1}$ | Death rate of actively infected cells. | Literature | $0.7 \text{ day}^{-1} = 21 \text{ month}^{-1}$, converting previous estimation from per day to per month <sup>85</sup>. |
| $\gamma$ | $1.05 \times 10^5 \text{ month}^{-1}$ | Production rate of virions. | Literature | Product of death rate $\mu_A$ (ref. <sup>85</sup>) and the viral burst size $n = 5000$ (ref. <sup>86</sup>). |
| $c$ | $1.5 \times 10^2 \text{ month}^{-1}$ | Clearance rate of free virions. | Literature | Rate converted to per month <sup>87</sup>. |
| $\lambda_f$ | $60 \text{ month}^{-1}$ | Inverse of the characteristic lifetime of antigens. | Literature | Converted to per month <sup>67</sup>. |
| $\gamma_f$ | $7.67 \text{ month}^{-1}$ | Strength of variability of antigenic environment. | Fitted | Adjusted to recover IUPM fluctuations consistent with observations <sup>88</sup>. |
| $\nu_L$ | $29.4 \text{ month}^{-1}$ | Division rate of latently infected cells. | Literature | Division rate of T cells, converted to per month <sup>67</sup>. |
| $\mu_L$ | $36.595 \text{ month}^{-1}$ | Death rate of latently infected cells. | Fitted | Adjusted to recover average decay in clones with low probabilities of reactivation <sup>88</sup>. |

**Supplementary Fig. 1.**
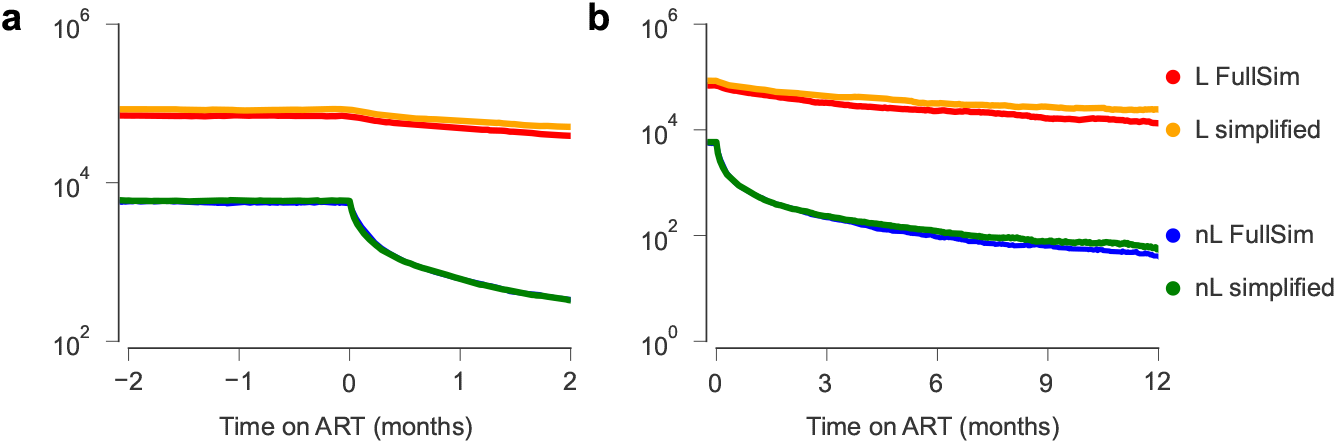
Comparison of decays between full simulation and simplified model. Decays of the total number of latently infected cells as well as the rate of new latent clones being produced. **a**, Start of ART. **b**, First year on ART.

**Supplementary Fig. 2.**
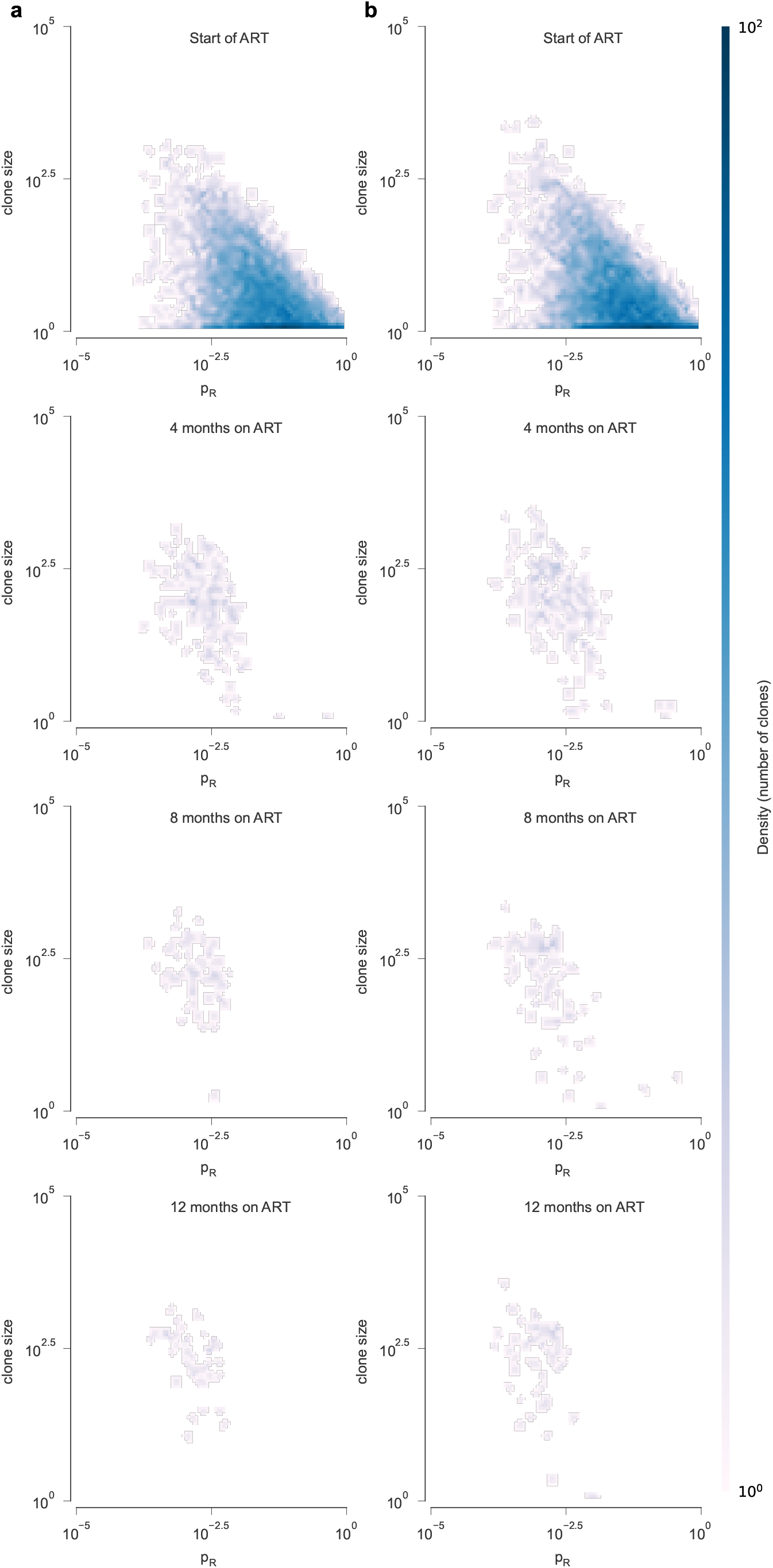
Comparison between full simulation and simplified model focusing on latently infected clones only. Side by side comparison of the clone size distribution in a *C* vs *p*_*R*_ plot between the full simulation (left) and the simplified one (right). Showing 4 timestamps during the first year on ART.

**Supplementary Fig. 3.**
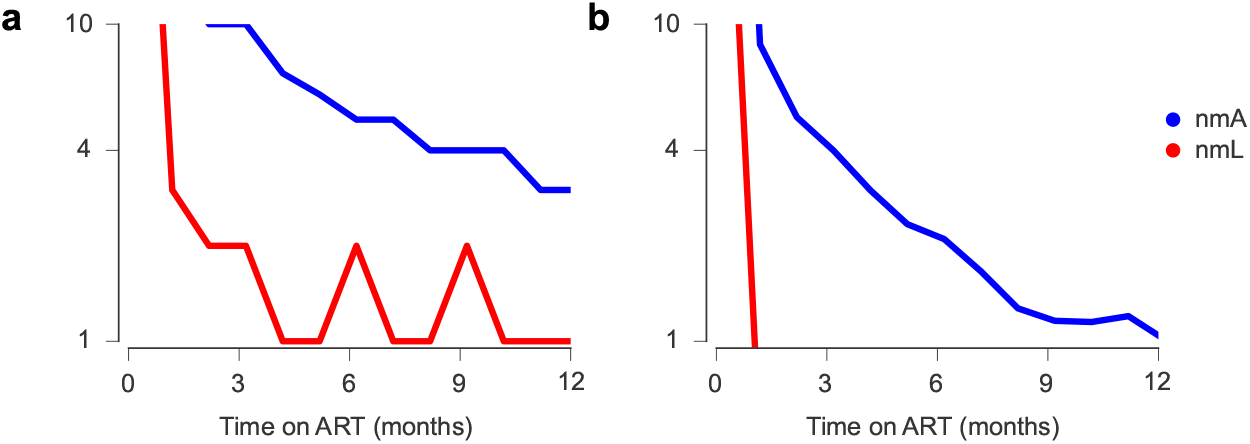
Max and mean number of new mutated sequences. Number of new cells with mutated pro-virus per time step. Upper corresponds to actively infected cells with a new mutated HIV sequence and the lower to new latently infected clones with new mutated HIV sequence. **a**, Max value over a window of a month. **b**, Average value over a window of a month.

